# Neural representations of the content and production of human vocalization

**DOI:** 10.1101/2022.09.30.510259

**Authors:** Vera A. Voigtlaender, Florian Sandhaeger, David J. Hawellek, Steffen R. Hage, Markus Siegel

**Affiliations:** Department of Neural Dynamics and Magnetoencephalography, Hertie Institute for Clinical Brain Research, University of Tübingen, Germany; Centre for Integrative Neuroscience, University of Tübingen, Germany; MEG Center, University of Tübingen, Germany; Graduate Training Centre of Neuroscience, International Max Planck Research School, University of Tübingen, Germany; Dept. of Otolaryngology-Head and Neck Surgery, Hearing Research Centre, University of Tübingen, Germany; Roche, Pharmaceutical Research and Early Development, Roche Innovation Center Basel, Basel, Switzerland

## Abstract

Speech, as the spoken form of language, is fundamental for human communication. The phenomenon of covert inner speech implies a functional independence of speech content and motor production. However, it remains unclear how a flexible mapping between speech content and production is achieved on the neural level. To address this, we recorded magnetoencephalography (MEG) in humans performing a rule-based vocalization task. On each trial, vocalization content (one of two vowels) and production form (overt or covert) were instructed independently. Using multivariate pattern analysis, we found robust neural information about vocalization content and production, mostly originating from speech areas of the left hemisphere. Production signals dynamically transformed upon presentation of the content cue, whereas content signals remained largely stable throughout the trial. In sum, our results show dissociable neural representations of vocalization content and production in the human brain and provide new insights into the neural dynamics underlying human vocalization.

## Introduction

Vocal behavior is an essential component of human communication. Particularly speech, the spoken form of language, is a highly sophisticated skill exclusive to humans. Thereby, we can encode information not only in sound (overt speech), but also in thought (covert speech). These different speech forms imply a functional independence of speech content and motor production. However, it remains an open question how content and production are represented neuronally and how the brain achieves a flexible mapping between the two.

In speech, two levels need to be distinguished. The lexical level, which refers to entire words, and the sub-lexical level, which refers to parts of words, such as phonemes or syllables ^1^. On the lexical level, neural activity underlying overt and covert speech is similar, with differences largely accounted for by a higher degree of executive motor control during overt speech ^2–5^. Broca’s area was found to act as a supra-modal hub, exhibiting language-specific activation independent of the production form ^6^. These findings suggest that there is a neural representation of speech content which is to some degree independent of its motor production. While this independence is intuitive on the lexical level, it is less clear on the sub-lexical level, where content may be expected to be tightly bound to the motor production. Still, initial evidence suggests such production independent representations of sub-lexical entities like syllables and phonemes ^7,8^.

In sum, past work suggests that neural activity underlying lexical and sub-lexical vocalizations represents both speech content and motor production. However, it remains unclear whether content representations generalize across different motor plans, and to what extent it is possible to dissociate these aspects. Furthermore, the dynamic interplay between emerging representations of content and motor production is not known, as most previous studies used neural data with inherently poor temporal resolution.

Here, we therefore aimed, first, to independently manipulate and decode neuronal content and motor components of human vocalization, and second, to investigate their dynamic interplay across time. We recorded MEG while subjects performed a novel rule-based vocalization task dissociating the content and motor aspects of sub-lexical speech. Content (one of two vowels) and production (overt or covert) were instructed sequentially and in random order. Multivariate pattern analysis (MVPA) of time-resolved MEG data allowed us to characterize the format, overlap and temporal dynamics of neural content and production representations.

With this joint approach, we were able to read out content and motor information several seconds before speech onset. The strength of neural information correlated with the degree of motor involvement and the two representations overlapped when isolated. The production representation transformed once the content was known, whereas the content representation remained stable until the onset of vocalization.

## Results

### The components of vocalization can be decoded independently

We recorded MEG while subjects performed a rule-based vocalization task. Participants had to overtly vocalize or covertly imagine the vocalization of two different vowels. During each trial, content (/u/ / /Ə/) and production (vocalized/ imagined) were instructed sequentially with visual cues (Fig. 1A). Each cue lasted 100 ms and was followed by a 2 s delay. At the end of the trial, a brief dimming of the fixation point served as a go-cue for the onset of vocalization or imagination. The order of instruction was randomized, as was the assignment of the instructed content or production to the visual cues (Fig. 1B). Participants performed the correct production type (vocalized vs. imagined) in 97.98 % of the trials. In case of vocalized trials, the correct vowel (/u/ vs /Ə/), was performed in 100 % of the trials. We checked vocalized trials for their onset latency (possible for 37 sessions). In 98.48 % of the trials, the vocal onset was after the go cue, as instructed. The mean vocal latency of these trials was 0.58 s (+/-0.12 SD).

**Fig. 1.**
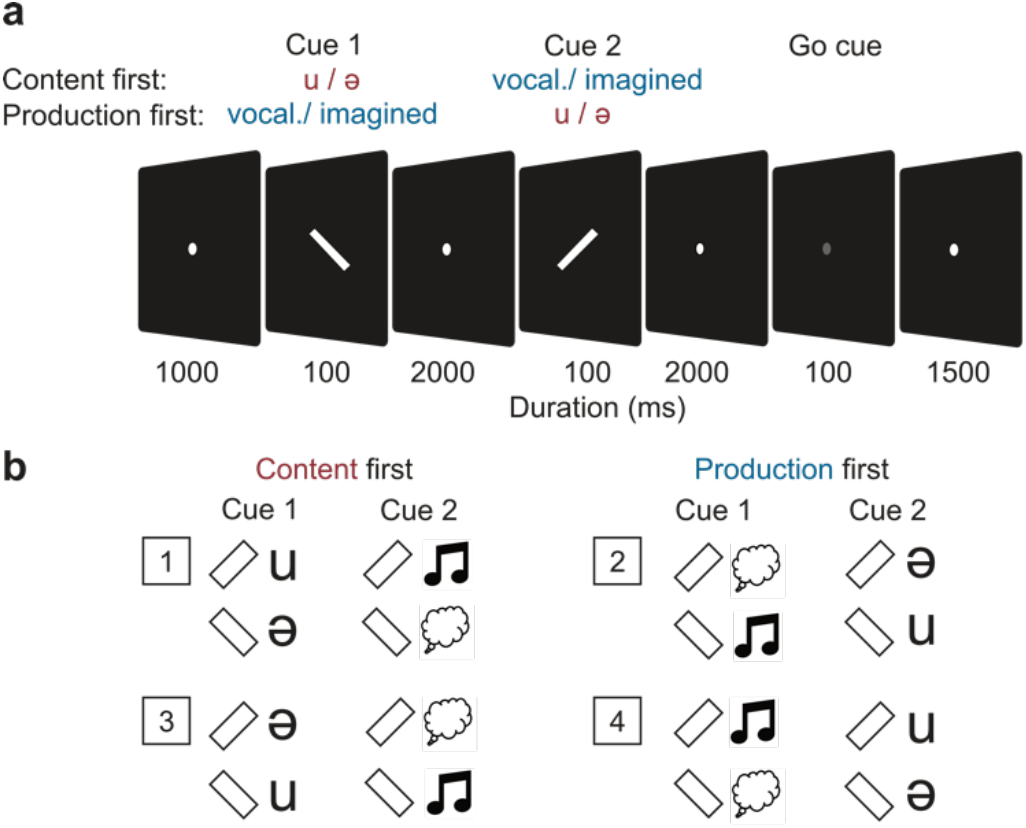
Rule-based vocalization task. Participants imagined or vocalized different contents (phonemes /u/ and /Ə/). **a**, Production and the content were instructed successively with visual cues, according to the respective rule of the trial block. **b**, Rules for four trial blocks per recording session. In two blocks the content was instructed first, in the other two blocks the production was instructed first.

For each subject and recording session, we computed neural information about content and production. We applied cross-validated MANOVA on preprocessed single-trial MEG data from all sensors (cvMANOVA; see Methods) ^9,10^. As an extension of the commonly used cross-validated Mahalanobis distance, cvMANOVA allows for the simultaneous quantification of the variability in neural data due to several variables of interest. The resulting measure of neural information can be interpreted analogously to classifier performance from multivariate decoding analyses. To enable robust statistical tests, we averaged information in the time windows between cue 1 and 2 (delay 1), as well as between cue 2 and the go cue (delay 2). In both cases, we excluded the first 250 ms after cue offset to avoid a possible confound due to sensory activity evoked by the cues (see Methods).

We observed significant neural information about both variables (Fig. 2). For both orders of cue presentation, we found information about the variables shortly after their respective instruction. When content was instructed first, there was significant content information in delay 1 (p = 0.003; corrected) and significant information about content and production in delay 2 (content: p = 3.4×10^−4^, production: p = 1.6×10^−9^; corrected). Conversely, when production was instructed first, content information was only present in delay 2, whereas production information was highly significant in both delays (content: p_del2_ = 7.8×10^−5^; production: p_del1_ = 2.8×10^−6^, p_del2_ = 8.5×10^−9^; corrected). Thus, both the content of a vocalization and its production form were represented neuronally, several seconds before the actual execution.

**Fig. 2.**
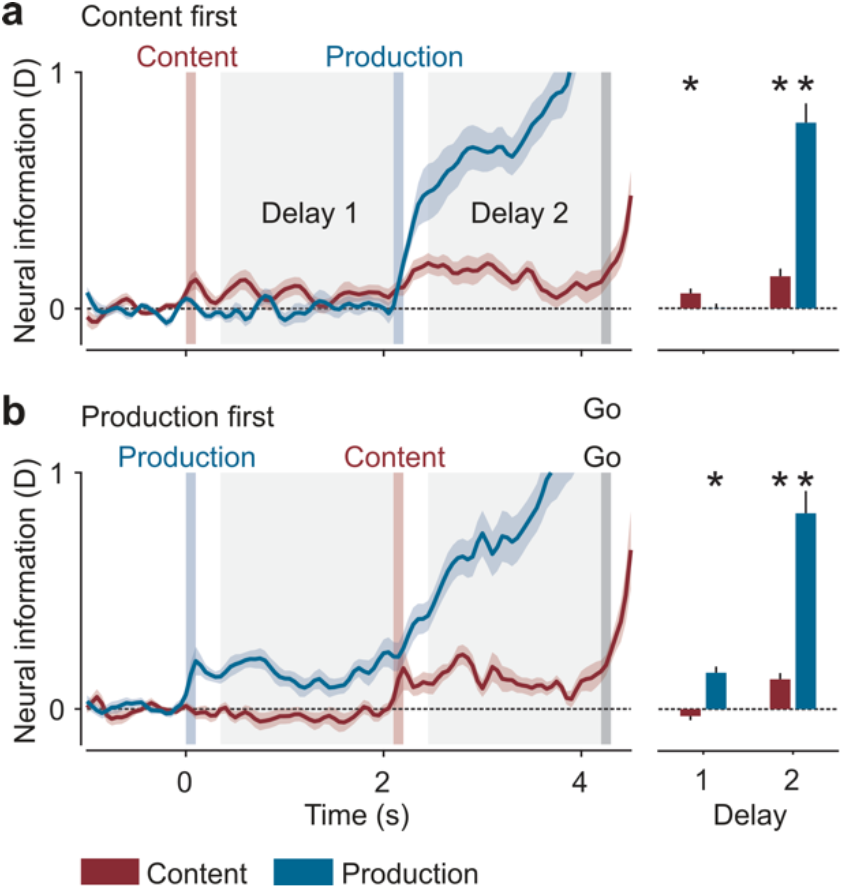
Neural information about content and production. **a**, Information in trials with content instructed first. **b**, Information in trials with production instructed first. Shaded regions and error bars indicate SEM. Bar plots show average information in delay 1 and 2. Asterisks indicate significance (n = 24, p < 0.05 corrected; t-test, one-tailed).

### The components of vocalization are modulated by effort

Both experimental dimensions entailed differences in motor effort. Imagined vowels lacked actual vocalization, just as /Ə/, as a non-articulated vowel, lacked the strong articulation of /u/. Therefore, we wanted to investigate whether the neural information about content was equally strong for both production types and whether the neural information about production type was identical for both vowels. To test this, we repeated the analysis after splitting the data based on both variables. In other words, content was decoded separately for both production types, and production was decoded separately for both vowels (Fig. 3).

**Fig. 3.**
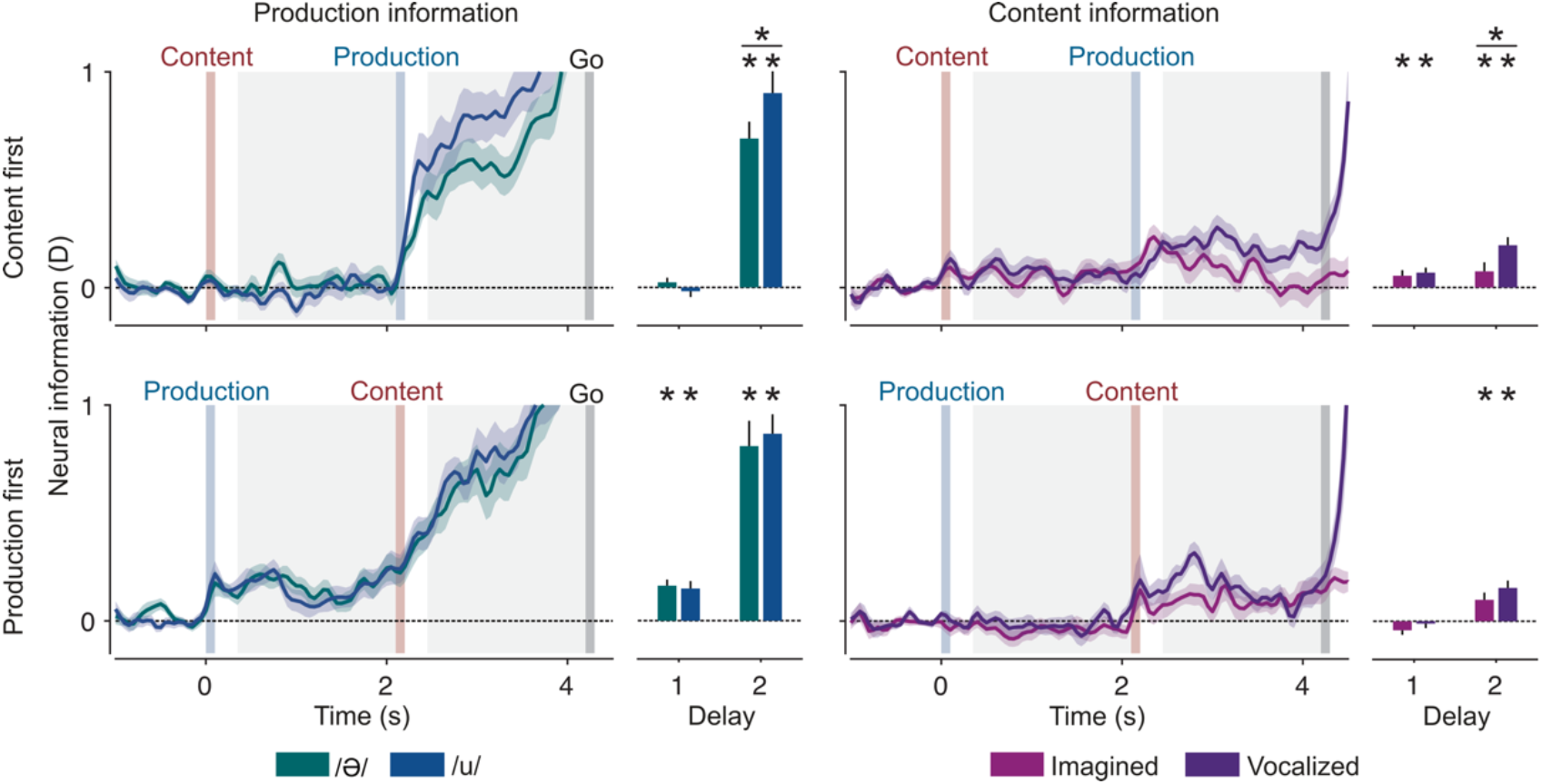
Neural information about content and production in split conditions. Shaded regions and error bars indicate SEM. Bar plots show averaged information of delay 1 and 2. Asterisks indicate above bars significant information (n = 24, p < 0.05 corrected; t-test, one-tailed). Horizontal lines with asterisks on top indicate a significant difference (n = 24, p < 0.05 corrected; paired t-test, two-tailed).

In all conditions, information about both variables could be decoded after their respective instruction. When content was instructed first, production information was significant for both vowels, in delay 2 (/Ə/: p = 7.8×10^−9^, /u/: p = 6×10^−9^; corrected). When production was instructed first, it could be read out in both delays and for both vowels (/Ə/: p_del1_ = 4.1×10^−6^, p_del2_ = 4.8×10^− 7^; /u/: p_del1_ = 1.4×10^−4^, p_del2_ = 1.6×10^−9^; corrected). In the second delay, production information was higher in /u/ than in /Ə/ trials, but only significantly so when content was instructed first (p = 0.026, paired t-test; corrected).

Content information was significant in both delays and production types when content was instructed first (imagined: p_del1_ = 0.044, p_del2_ = 0.044; vocalized: p_del1_ = 0.004, p_del2_ = 2.7×10^−5^; corrected). When production was instructed first, content information was only significant in the second delay, again in both production types (imagined: p = 0.007, vocalized: p = 1.5×10^−4^; corrected). Content information was higher during vocalized than imagined trials, in both orders and all relevant delays. This difference was significant in the second delay when content was instructed first (p = 0.007, paired t-test; corrected).

In sum, both production and content information were present in all individual conditions and, furthermore, higher in those conditions with stronger motor involvement.

### The components of vocalization are represented in cortical speech areas

We found that vocalization content and production were represented in cortical areas typically associated with speech. To characterize the cortical distribution of content- and production information, we repeated the cvMANOVA-analysis on the source level using a searchlight approach. We then averaged neural information within four 500ms time windows per delay. Again, for the 1st time window after each cue, the initial 250ms were excluded. This analysis revealed spatially stable representations of both variables (Fig. 4). Descriptively, the highest level of production information was found in Broca’s area and other motor cortices. Content was represented in the same areas, but additionally in temporal cortices.

**Fig. 4.**
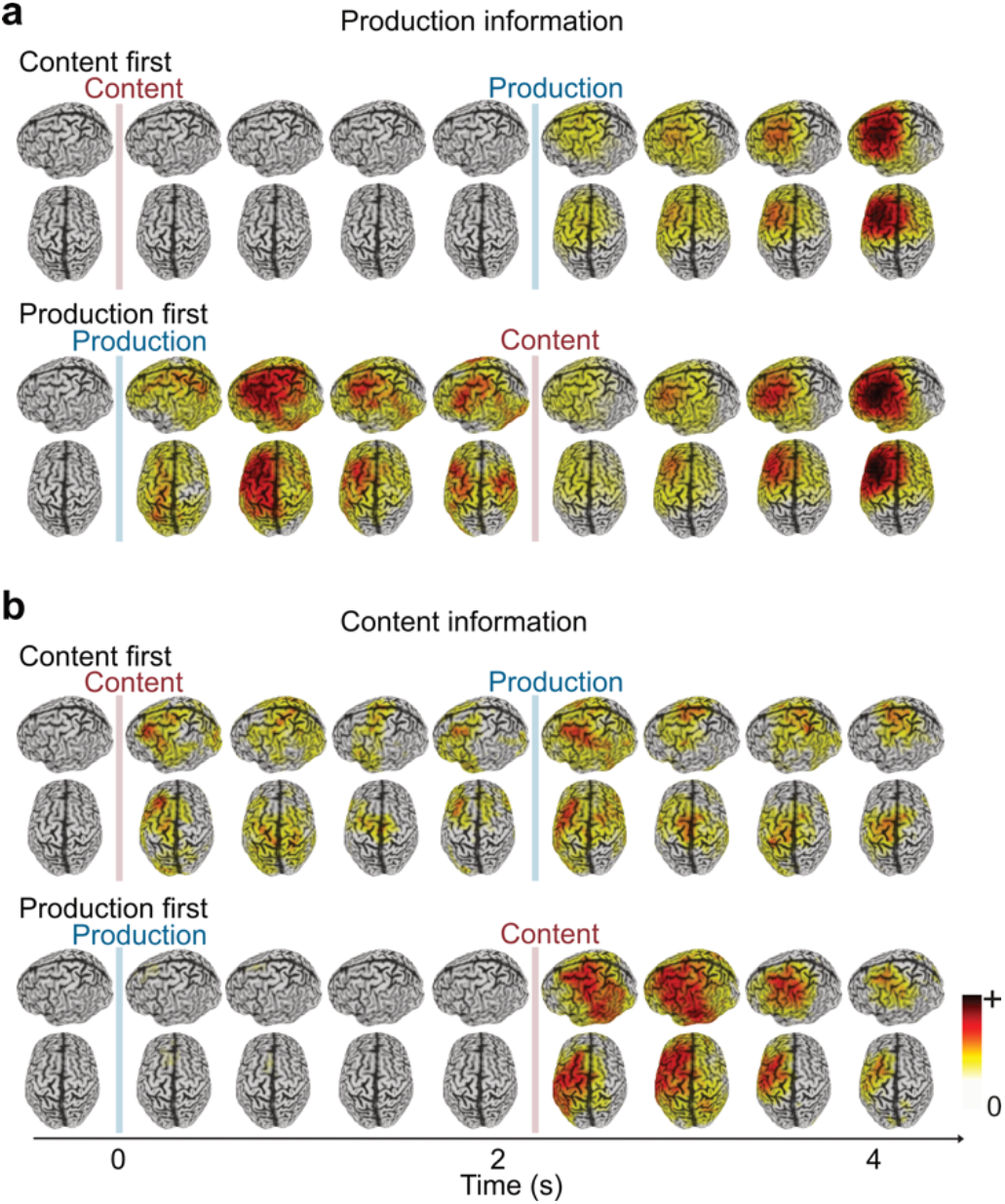
Spatial dynamics of neural information about content production. **a**, Production information in both orders. **b**, Content information in both orders. Left and top view of the cortical distribution of information. Information was averaged in 500 ms intervals, except for the intervals directly after the cues, for which the first 250 ms were excluded.

For higher-order language processes neural activity is known to be left-lateralized, which is debated for speech on the sub-lexical level ^11,12^. Our source patterns suggested a clear left-lateralization. We tested this by computing a lateralization index for both variables (information in the left hemisphere -right hemisphere, Fig. 5). Both variables were left-lateralized in all delays after the respective instruction. This was significant for content information in the first delay when it was instructed first (p = 0.037; corrected) and in the second delay when production was instructed first (p = 0.006; corrected). Production information was significantly left-lateralized in both delays when production was instructed first (p_del1_ = 0.048, p_del2_ = 0.009; corrected) and in the second delay when it was instructed second (p = 0.003; corrected).

**Fig. 5.**
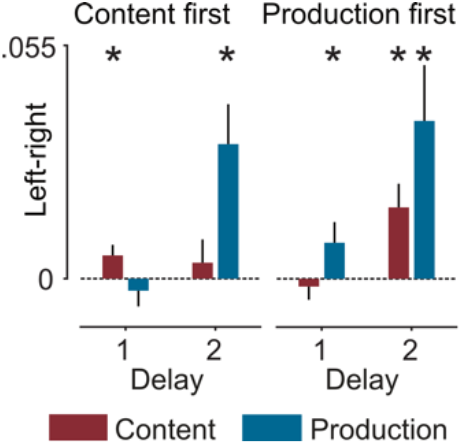
Lateralization index of content and production information. The lateralization index (left – right hemisphere information) was computed for both delays and both orders. Asterisks indicate significance (n = 24, p < 0.05 corrected; t-test, one-tailed).

Taken together, source analysis showed that both content and production had stable left-lateralized representations on the cortical level. This did not only identify the origin of neural information within well-known speech-associated areas, but also excluded confounds due to inherently non-lateralized effects of visual cues or electromagnetic artifacts.

### The components of vocalization have different representational formats

The searchlight analysis indicated a spatial stability of the coarse cortical distribution of content and production information across time. Nonetheless, the representational format, i.e. the fine-grained cortical pattern underlying each representation may be dynamic. To test if neural representations of content and production transformed across time, we decoded both variables on the sensor level across time ^13^ (Fig. 6). By using each pair of time-points for training and testing the cvMANOVA, we could assess to what extent neural representations were overlapping across timepoints. Stable or dynamic representations would yield high or low cross-time decoding, respectively. We performed this analysis both on the content-first condition and on the production-first condition, as well as training on all timepoints of one condition and testing on those of the other (Fig. 6, mixed orders).

**Fig. 6.**
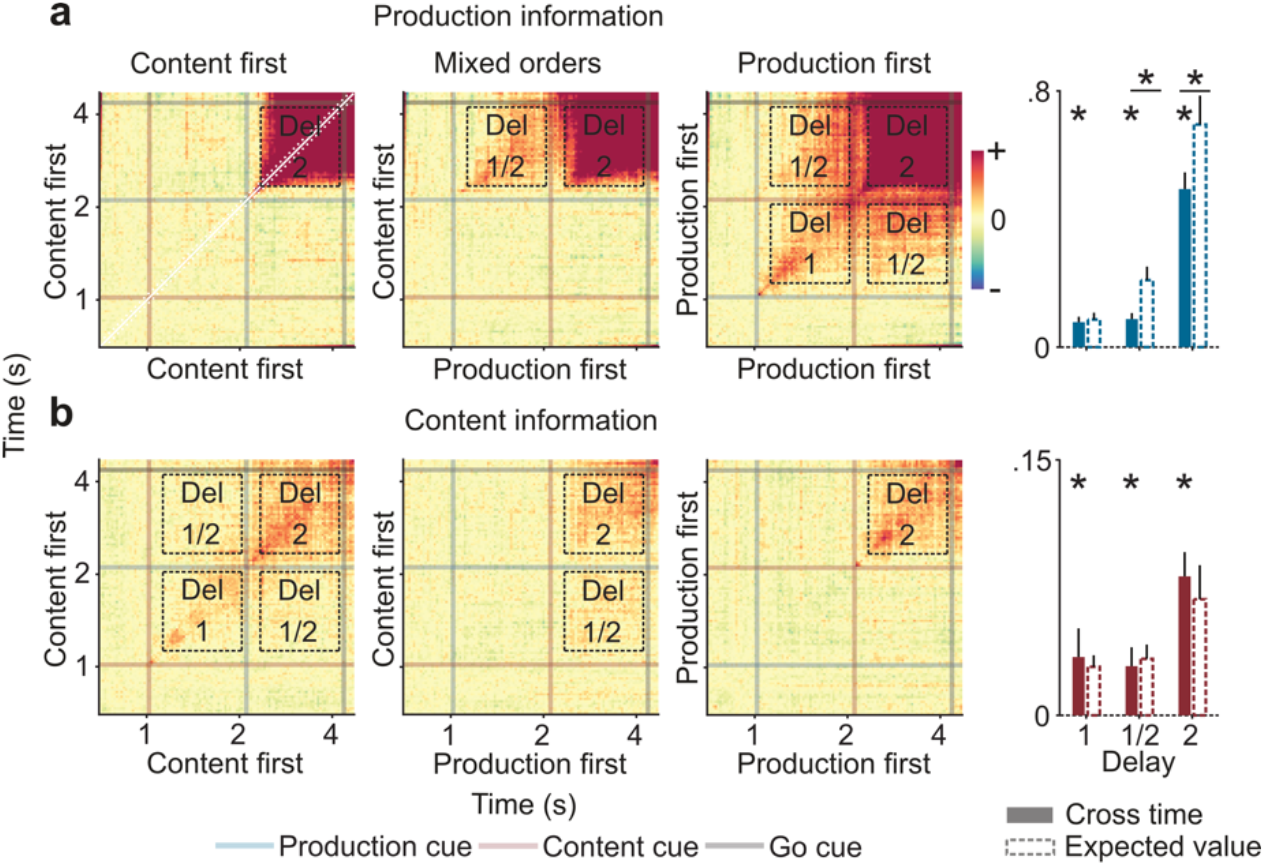
Cross-time information about content and production. **a**, Cross-time production information. **b**, Cross-time content information. Bar plots show the averaged cross-temporal information for delay 1, 2 and 1/2, as well as the expected values for stable representations. The diagonals ± 100 ms were excluded from calculations. Asterisks indicate significant information (n = 24, p < 0.05 corrected; t-test, one-tailed) and horizontal bars with asterisks on top indicate significantly smaller cross-information than the expected value (n = 24, p < 0.05 corrected; paired t-test, one-tailed).

For statistical testing, we averaged neural cross-information per variable and delay (Fig. 6, bar plots). Cross-temporal Information within delay 1 was averaged over all conditions in which the relevant cue was instructed first, as information could only be present in these conditions. To estimate cross-temporal generalization between delays, we averaged an off-diagonal time-window over those conditions where the relevant cue was presented first, as well as those mixed-order conditions where the relevant cue was presented first in one condition but second in the other. To estimate cross-temporal information within delay 2, we averaged data from all cue orders. Briefly, this ensured that neural information about content or production was available at all training and test timepoints included in the statistical analysis, allowing us to meaningfully address the question whether representations were shared between training and test timepoints.

Within both delay 1 and 2, there was significant cross-temporal information about both variables (production: p_del1_ = 6.3×10^−5^, p_del2_ = 2.9×10^−9^; content: p_del1_ = 0.027, p_del2_ = 1.1×10^−5^; corrected). There was also significant cross-temporal information between delays 1 and 2 for both variables (production: p = 6.3×10^−5^; content: p = 0.012; corrected).

To quantitatively interpret these results in relation to the information available at each individual timepoint, we computed an estimate of the expected cross-information in case of perfectly stable representational formats but potentially different information magnitudes over time ^10^. This expected cross-information allowed us to test whether representations significantly differed between timepoints. The cross-temporal stability of production representations was lower than expected in all time windows, and significantly so between delay 1 and 2 and within delay 2 (p_del1/2_ = 2×10^−4^, p_del2_ = 7.3×10^−6^; corrected). In contrast, cross-temporal content information was never significantly smaller than expected for a temporally stable representation. Thus, while we found evidence for a partially dynamic representation of production type, this was not the case for content, which appeared stable over time.

To what extent are the neural representations of content and production similar? Our cross-time decoding analysis showed that both representations evolve differently across time. We took this as an indication that these representations are not identical. To rigorously estimate the extent of representational overlap, we implemented a cross-variable analysis, training the algorithm on one variable and testing it on the other. Again, we computed the expected cross-information under the assumption of identical representations and compared it with the observed cross-information. We computed cross-variable information in both cue orders, but also in mixed cue orders where the relevant cue was either first or second (Fig. 7).

**Fig. 7.**
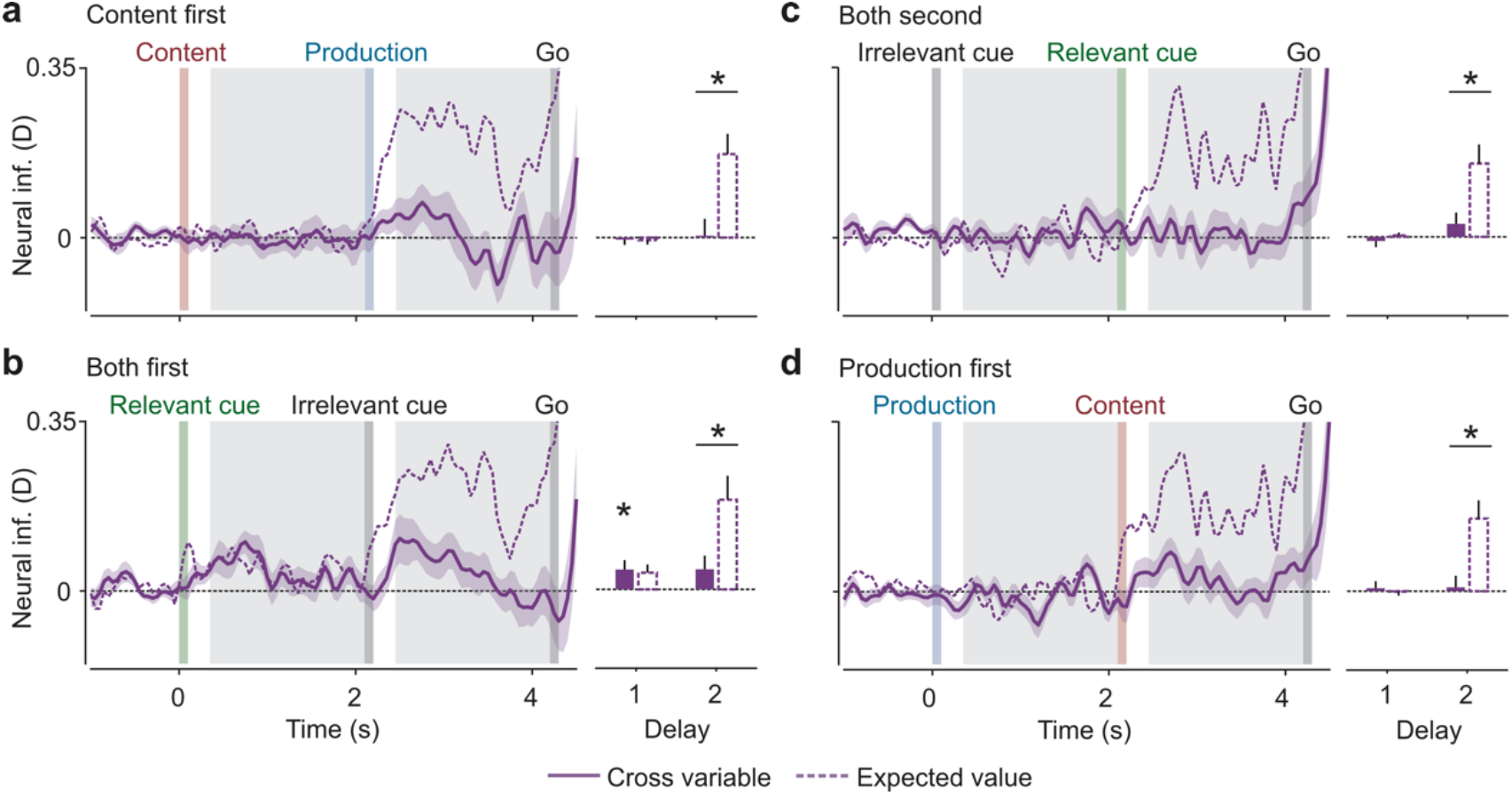
Cross-variable information of content and production. Cross-information in both orders and mixed orders with the relevant cue first or second. **a**, Cross-information in trials with content instructed first. **b**, Cross-information across trials with relevant cue first. **c**, Cross-information across trials with relevant cue second. **d**, Cross-information in trials with production instructed first. Solid lines indicate cross-information and dashed lines indicate the respective expected values for identical representations. Bar plots show the averaged cross-information and the averaged expected values. Asterisks indicate significant cross-information (n = 24, p < 0.05 corrected; t-test, one-tailed) and horizontal bars with asterisks on top indicate significantly smaller cross-information than the expected value (n = 24, p < 0.05 corrected; paired t-test, one-tailed).

**Fig. 8.**
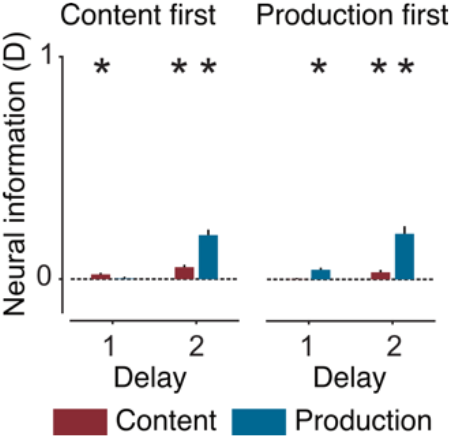
Cross-session decoding of content and production. Information was averaged for both orders and delays. Asterisks indicate significant information (n = 24, p < 0.05 corrected; t-test, one-tailed).

We found significant cross-information only in delay 1 in the order with the relevant cue first (p = 0.046; corrected). In the other orders, as information about both variables could only be present after the second cue, no cross-information was expected in delay 1. While we observed a small amount of cross-information in delay 2 of the mixed orders, this was not significant. On the other hand, cross-information was significantly smaller than its expected value in the second delay of all orders (content first: p = 0.004; both first: p = 0.02; both second: p = 0.01; production first: p = 0.002; corrected). Taken together, these results show that content and production representations overlap but are not identical. In our data, isolated content representations (before knowledge of the production type) cannot be distinguished from isolated production representations (before knowledge of the content). Thus, content and production representations are indistinguishable as long as the respective other variable is unknown to the participant. However, as soon as both aspects become available, their representations show strong differences.

### Information about the components of vocalization remains stable across sessions

Electromagnetic artifacts and other, usually visually driven confounders, often pollute the data in speech studies ^14^. Our source level analysis provided evidence that content- and production information was indeed speech related, originating from well-known speech-associated areas. In addition, we implemented a control analysis to test if a possible visual cue confound had an impact on our results. Because the order of trial blocks was reversed in each participant’s second recording session, it was possible to cancel out the visual cue effect by decoding across sessions. To this end, we trained the cvMANOVA on one session and tested on the other.

Decoding across sessions was expected to be more challenging than decoding within session, as the signal-to-noise ratio was impacted by additional variability due to head movement between the sessions. Nevertheless, we found significant information about both content and production in all relevant delays. There was significant content information in both delays if it was instructed first (p_del1_ = 0.004, p_del2_ = 5.1×10^−5^; corrected) and in delay 2 if it was instructed second (p = 0.006; corrected). Production information was significant in delay 2 if content was instructed first (p = 3.4×10^−8^; corrected) and in both delays if production was instructed first (p_del1_ = 4.8×10^−5^, p_del2_ = 3.4×10^−6^; corrected). Thus, content- and production information were stable across recording sessions, and therefore also not driven by a visual, cue-related confound due to the sequential order of rule blocks within each session.

## Discussion

Our results shed new light on the neural mechanisms underlying the flexible mapping between two components of human speech: its content and motor production. Combining MEG and MVPA, we could decode robust neural information about these components directly after their respective instruction, which was up to several seconds preceding the onset of the vocalization. Thus, our factorial task design allowed us to dissociate content and production in the pre-execution phase, where key processing stages take place ^15–18^ and electromagnetic artifacts of the motor production itself are ruled out ^14^.

We found significant information about each component directly after the first cue, when the respective other instruction was still missing. The early content information suggests that content can be represented independently of a specific motor plan, which falls in line with implications from previous studies ^2,6–8,12,15,19^. Therefore, the actual motor production is not necessary to form a neural representation of content. However, this does not imply that the content representation is completely independent from motor planning.

Content information was present for both production types, as was production information for both vowels. However, content information was higher when vocalized and production information was higher for the vowel /u/, implying a dependence of information strength on the degree of motor involvement. As previous studies have shown, overt and covert speech differ in terms of executive motor control, including M1 recruitment for the efference copy ^2,5^, which could account for the higher content information in vocalized trials. In addition, the stronger phonological code retrieval and encoding in overt speech could also contribute to the observed difference of content information between the production types ^5^. The two vowels also differ in motor involvement, as /Ə/ is a non-articulated innate-like vowel, whereas /u/ is strongly articulated and learned ^20^. Therefore, the stronger motor involvement could account for higher production information in the vowel /u/.

The vocal motor network comprises many cortical and subcortical areas (Hage and Nieder, 2016; Jürgens, 2002; Schulz et al., 2005). As expected, we located representations of content and production in frontal and central cortex consistent with well-known speech-associated areas. In addition, content representation extended to temporal cortices, which may be due to efference copies in sensory regions ^23–25^. Although one may not expect the higher-order language network to be recruited in our paradigm, we found neural representations of both content and production to be stronger in the left hemisphere. This could either indicate that language capacities beyond low-level speech were recruited or that low-level speech processes can already be lateralized under specific circumstances. Independent of these alternatives, our finding of lateralization in an early pre-execution phase suggests that the uncovered neural representations were indeed speech or language specific and did not reflect general working memory processes.

Our cross-decoding approach provided new insights into the temporal dynamics and similarities between the neural representations of vocalization components. During the first delay, when information about only one of the components was available to the subjects, both representations were temporally stable and indistinguishable from each other. Thus, while there may be subtle differences that MEG is insensitive to, our results suggest an overlap of neural representations of content and production. One possible explanation for such an overlap could be an effort effect where conditions with a higher degree of motor involvement elicit higher neural activity than those with a lesser degree. Concretely, |u vs. Ə| and |vocalized vs. imagined| could both correspond to contrasts of |high effort vs. low effort|. This effect could reflect priming motor signals preceding vocalizations ^18,26^. Alternatively, it could also reflect the firing patterns of one or more speech specific neural populations that encode several content- and production-related features. Here, effort could drive either the firing rates of individual neurons or the number of recruited cells. Further invasive research is required to determine whether the same population, or spatially close and therefore indistinguishable neurons are modulated by both content- and production-related effort.

In the second delay, cross-information between content and production was similar as in the first delay. However, as both content- and production information were much higher, the representations were now clearly distinguishable. Moreover, cross-temporal decoding between both delays revealed that the content representation remained stable over time, whereas the production representation transformed once the content was known. Consequently, the divergence of the representations in the second delay was likely driven by the transformation of production. Taken together, this implies that the production representation during the second delay was a combination of the initial format during the first delay and an additional component. This additional component, building up once the content was known, may reflect the specific motor program used to prepare the articulation of the respective vowel.

In sum, our results show that, when isolated, both representations overlap and correlate with the degree of motor involvement. This overlap of the representations could be caused by pre-motor-like activity that is modulated by effort, but independent of the actual execution. While the content representation remains stable, the production representation changes once the content is known, likely reflecting the addition of a specific motor program.

While natural speech can functionally be decomposed into content and motor production, there is little evidence for a content dimension on the neural level and even less about its dynamic interplay with the motor production. Most implications for a neural content dimension come from studies focusing on the lexical ^3,6,24^ or the sub lexical syllable level ^2,7,23^. Temporal dynamics have so far only been studied on the lexical level ^15,16^. Yet, the elementary building blocks of speech are phonemes, and to our knowledge all previous research on this level is related to motor production ^12,19,27,28^. Our results uncover a neural content dimension for phonemes that was present independently of motor production and could therefore allow for a generalization between production forms. These results accord well with the larger body of work on the higher sub lexical and lexical levels.

Our findings set the stage for future research, to investigate how neural codes of isolated phonemes and their motor production translate to those embedded in speech. Our combined approach of MEG, MVPA and a rule-based paradigm, provides a fruitful framework for that, thus, opening a new window for non-invasive speech research in health and disease.

## Methods

### Subjects

24 healthy humans with normal or corrected-to-normal vision participated in the study (14 male; 21 right-handed; mean age: 29 years; 5 years SD). All participants gave written informed consent before participation and received monetary reward afterwards. The study was conducted in accordance with the declaration of Helsinki and approved by the ethics committee of the University of Tübingen.

### Behavioral Task and Stimuli

Participants performed a rule-based vocalization task. In each trial, one of two vowels (/u/ or /Ə/ had to be either overtly or covertly vocalized. Vowel and production type were instructed sequentially by visual cues. The corresponding rule, i.e. the assignment of the visual cues to their instructed content, changed across recording blocks and was indicated before the beginning of each block.

Participants self-paced the trials using closed-loop eye movement control. Each trial started with an initiation phase of 1000 ms, during which a white fixation spot (diameter: 0.1° of the visual angle) appeared at the center of the screen. Once fixation was acquired, the first visual cue (a forward or backward white slash, length: 2°, width: 0.25° of the visual angle) appeared for 100 ms, instructing either content or production. Then, the fixation spot appeared again for a delay period of 2000 ms. The second visual cue appeared for 100 ms, instructing the respective missing variable, followed by a second delay period displaying the fixation spot. Dimming the spot for 100 ms served as the go-cue for the participants’ response. After the go-cue, the fixation spot remained on screen for 1500 ms, which gave time for the response of the participants. The intertrial interval was 1000 ms long, indicated by dimming of the fixation spot. If fixation was broken after the onset of the first visual cue, the trial was aborted, which was indicated by a color change of the fixation spot to red for 500 ms. The cue configuration of the aborted trial was repeated at a random position later within this block.

Within each recording block, the order of instruction and the meaning of each instruction cue was fixed. However, in half of the blocks, the content was instructed first, whereas in the other half of the blocks, the production was instructed first. Moreover, the assignment of each visual cue to its meaning (content or production) was different in half of the blocks for each order. These four different rules led to four blocks of trials. Each block contained 80 trials, with 20 per condition (1: /u/ vocalized, 2: /u/ imagined, 3: /Ə*/* vocalized, 4: /Ə/ imagined). The order of the conditions was randomized per block. In total, there were 16 different conditions, including the different assignments of the visual cues to their instructed content. The 24 possible orders of the four blocks were randomly assigned to the participants.

Before the experiment, participants were instructed on how to articulate the vowels correctly. Thereby, we made sure that /u/ was clearly articulated, whereas /Ə/ was produced with minimal involvement of the vocal tract. We also made sure that there was no involuntary articulatory movement visible for imagined vowels. During the experiment, the participants memorized the respective rule before each block. The rule was presented on the screen for as long as necessary. Eight training trials preceded each block to ensure correct performance. If necessary, the rule was shown again during the block while the sequence of trials was paused. All participants performed two MEG sessions with 320 trials each. For each participant, the order of blocks from the first session was inversed for the second session. During both sessions, the performance of the participants was monitored with a microphone and a camera. After the experiments, each trial was labeled for production type and in case of vocalized trials for vowels.

### Data Acquisition

We recorded MEG (Omega 2000, CTF Systems Inc., Port Coquitlam, Canada) with 273 sensors at a sampling rate of 2342.75 Hz in a magnetically shielded chamber. Participants sat upright with a screen at 65 cm viewing distance. Stimuli were projected onto the screen by an LCD projector (Sanyo PLC-XP41, Moriguchi, Japan) with a refresh rate of 60 Hz. The projection was the only source of light in the chamber. Continuous head movement was monitored with three coils attached to fiducial points. In two participants, head movement could not be measured due to technical issues. Eye movements were recorded using an infrared eye-tracker (EyeLink CL-OC, SR Research Ltd., Canada) at a sampling rate of 1000 Hz. For labeling of trials and vocal onset detection in vocalized trials, the participants’ responses were recorded with a microphone integrated into the MEG System, at a sampling rate of 2343.75 Hz. An additional microphone (MD 419, Sennheiser electronic GmbH & Co KG, Germany) with a sampling rate of 44.1 kHz was used to record a higher quality audio trace for the labelling of production types and vowels. In a separate session, we acquired structural T1-weighted MRIs (3 Tesla MAGNETOM, Siemens Healthcare GmbH, Germany) for source reconstruction based on each participants individual’s anatomy (resolution: 1 mm^3^, MPRAGE).

### Data Preprocessing

Technically caused channel jumps were detected and corrected and time lags between digital triggers and actual stimulus presentation were corrected based on a photodiode signal. For one subject, we excluded a noisy channel from the analysis. We low-pass-filtered the MEG data at 30 Hz (6^th^ order, zero-phase Butterworth IIR filter) and down-sampled to 300 Hz. Each trial was baseline-corrected using the 500 ms preceding the onset of the first visual cue. To reduce electromagnetic artifacts, we ran independent component analysis (ICA) on the data. To ensure convergence of the ICA algorithm, the data was high-pass-filtered at 0.05 Hz. However, we applied the resulting unmixing matrix to the original data without any high-pass filter, and removed artefact components like heart beats and other small muscle activities. Vocal onset detection in vocalized trials was possible for 37 sessions. Audio traces were missing due to technical issues in the remaining session. However, performance was closely monitored during the recordings and immediately corrected, if participants vocalized before the go-cue appeared. The available audio traces were smoothed with a median-based sliding window model (window size 42.66ms). A participant-wise threshold served for onset detection. For that, the root mean square (rms) of a 1000 ms period of one of the imagined trials was calculated. The respective standard deviation was multiplied by eight and added to the rms. In case of low vocalization amplitudes due to very soft voices of a few participants, the threshold was manually adjusted by slightly decreasing it until the onsets could reliably be detected. The vocalization was correctly performed, with an onset after the go-cue, in 99% of the trials.

For all subsequent analyses, only correct trials were used. Those were trials with the correct production type, and, in case of vocalized trials, with the correct vowel and a vocal onset after the go-cue. For the nine sessions with a missing audio trace, only the correct vowel was considered.

### Cross-validated MANOVA

We estimated the amount of neural information about the variables of interest in the MEG data with cross-validated MANOVA ^9,10^. We performed 20 repetitions of cvMANOVA for each session with five-fold cross-validation. All folds and repetitions were subsequently averaged. We first estimated a noise covariance matrix using trials from all conditions. Next, we estimated contrasts of beta weights of each condition in a cross-validation fold’s training set, which accounted for the “training” of the model. The “testing” was done by estimating contrasts of beta weights in the respective fold’s test set. The dot product of these contrasts, normalized by the noise covariance, served as an estimate of the true pattern distinctness:

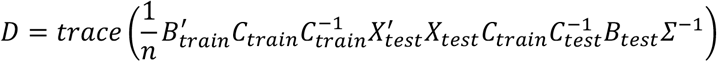

where Σ^−1^ is the inverted noise covariance matrix, C_train_ is the contrast vector the model is trained on, C_test_ is the test contrast vector and X_test_ is the design matrix indicating the unique condition of each trial in the test set. B_train_ and B_test_ contain the regression parameters of a multivariate general linear model:

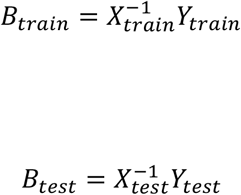

where Y_train_ and Y_test_ are the training and test data sets. The inverted noise covariance matrix was estimated with the mean of the time window from cue 1 offset to go cue onset:

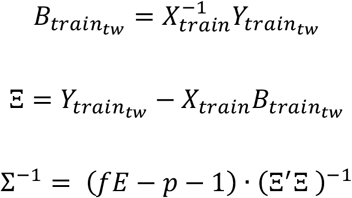

with fE as the degrees of freedom and p as the number of sources.

Technically, cvMANOVA is a multivariate information based cross-validated encoding approach. However, it shares many similarities with common multivariate decoding methods ^29^. The measure of neural information about the variables of interest can, theoretically, also be used to decode these variables on individual trials. Therefore, we refer to our results as decoding results.

For all sensor level analyses, a subset of 137 approximately equally distributed sensors were included. This was to ensure a sufficient number of trials in relation to the degrees of freedom of the dataset.

### Cross-decoding

With cvMANOVA we were able to decode across conditions, by training and testing on different time points, variables, and levels of the variables. Therefore, the contrast vectors C_train_ and C_test_ were constructed to only contain the respective conditions to be trained or tested on. With this approach, we decoded across variables by using content (/u/ vs. /∂/) for the construction of the training contrast C_train_ and production type (vocalized vs. imagined) for the construction of the test contrast C_test_. To estimate content information for both production types separately, we constructed C_train_ based on trials including both production types, but C_test_ based on trials with only one production type, respectively. The same principle was applied for separately decoding the production type from both vowels. For decoding across time, we used the regression parameters B_train_ from one time point and B_test_ from another. We applied this to all pairs of time points. For cross-session decoding, we implemented a two-fold cross-validation such that trials from session 1 and 2 served as training and test sets alternately.

### Expected cross-decoding

To estimate a benchmark for the overlap between representations, we computed the expected cross-information ^10^. As the maximally possible amount of shared information between contexts depends on the information available in each individual context, the strength of the shared representation must be compared to the strength of both representations. Therefore, we estimated the expected cross-decoding

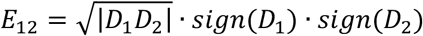

where D_1_ and D_2_ denote the pattern distinctness in the two contexts. If the representations were identical, the cross-decoding D_12_ would approach E_12_. Conclusively, cross-decoding values smaller than E_12_ indicate that the representations are not identical and, therefore, not fully overlapping.

### Source estimation

We generated individual single-shell head models ^30^ based on each subjects’ structural T1-weighted MRI. Using linear spatial filtering ^31^, we estimated three-dimensional MEG source activity at 457 equally spaced locations ∼7 mm beneath the skull. For searchlight analysis, we used the three dipole directions of each source and the respective immediate neighbors. The lateralization index (LI) was computed by averaging the searchlight results for each hemisphere and subtracting right from left. For cross-session decoding, the three orientations were added and a subset of 229 equally spaced sources was used for decoding.

### Statistical Analysis

Neural information was averaged within two different time windows (delay 1: from 250ms after cue 1 offset to cue 2 onset; delay 2: from 250ms after cue 2 offset to go-cue onset). For testing the significance of neural information and cross-information to be larger than 0 we employed one-tailed one-sample t-tests. One-tailed paired t-tests were applied for testing cross-time and cross-variable information to be smaller than the expected cross-information. For the comparison of content information in both production types and vice versa, we used two-tailed paired t-tests. All p-values were FDR corrected for the number of tested time intervals ^32^.

### Potential cue confound

We identified the source of a potential confound: due to the blocked design, in combination with possible non-stationarities in the dataset, content- and production information could theoretically be influenced by representations of the physical cue itself, even though the cue appearance and its meaning were counterbalanced. Briefly, if noise led to independent shifts of the neural activity patterns of both cue options, this could mistakenly be identified as information about the variable indicated by the cue. To make sure our results were not dependent on this potential confound, we took two measures. First, we excluded the 250 ms after cue onset in both time windows used for statistical analysis, as this was the time period that would likely be affected. Secondly, we performed cross-session decoding to confirm that content- and production information were present if the possible cue confound was accounted for.

### Visualization

For all line plots, data were smoothed with a 100 ms Hanning window (full width at half maximum).

### Software

All analyses were performed using the Fieldtrip toolbox ^33^ and custom code in MATLAB (R2014a & 2017b).

## Data availability

The preprocessed data are available from the authors upon reasonable request.

## Code availability

The analysis code is available from the authors upon reasonable request.

## Acknowledgements

We thank Gabriele Walker-Dietrich, Jürgen Dax and Christoph Braun for help with data acquisition. This research was supported by the European Research Council (ERC) StG 335880 (M.S.) and CoG 864491 (M.S.) and by the Evangelisches Studienwerk Villigst (V.V.).

## Author Contributions

Conceptualization: M.S., V.A.V., F.S., D.H., S.R.H.; investigation: V.A.V.; formal analysis: V.A.V., F.S.; resources: F.S., D.H., M.S.; writing – original draft preparation: V.A.V.; writing – review and editing: F.S., M.S.; supervision: M.S., F.S.; funding acquisition: M.S., V.A.V.

